# In vivo hemostatic capability of a novel Tetra-PEG hydrogel

**DOI:** 10.1101/2021.08.18.456884

**Authors:** Shinya Okata, Katsuyuki Hoshina, Kazumasa Hanada, Hiroyuki Kamata, Ayano Fujisawa, Yuki Yoshikawa, Takamasa Sakai

**Affiliations:** Department of Vascular Surgery, The University of Tokyo, Tokyo, Japan; Department of Bioengineering, The University of Tokyo, Tokyo, Japan

**Author notes:** **Corresponding author:** (KH).

## Abstract

TetraStat is a novel synthetic sealant created with a tetra-armed polyethylene glycol (PEG) hydrogel. It has no risk of infection from biological pathogens and has a hemostatic mechanism independent of the blood coagulation pathway and controllable gelation. We evaluated the hemostatic effect of TetraStat in ex vivo and in vivo experiments for future clinical application. In ex vivo experiments using a circulatory system filled with phosphate-buffered saline under high pressure, needle punctures were astricted with TetraStat and two commercially available hemostatic agents (SURGICEL and TachoSil). For in vivo experiments, rat vena cavae were punctured with 14, 18, and 20 gauge needles, and hemorrhage occurred for several seconds. A porous PEG sponge soaked with TetraStat was applied as a hemostatic system for the massive hemorrhage. In the ex vivo experiment, punctures were sealed completely after 1 min astriction with TetraStat gel; in contrast, SURGICEL and TachoSil failed to seal the hole. In vivo experiments demonstrated that TetraStat successfully caused hemostasis in the punctured vena cava within 1 min of application in a dose-dependent manner. For SURGICEL and TachoSil, successful hemostasis occurred after 5 min astriction but was less frequent after 1 min astriction. Ex vivo and in vivo experiments revealed TetraStat’s high hemostatic ability under high pressure and in rat vena cava injuries under massive hemorrhage. A porous PEG sponge soaked with TetraStat is a promising advancement in hemostatic systems.

## Introduction

Hemorrhage control is crucial for successful surgical procedures. Even after attempting to control hemorrhage by suturing the bleeding sites, surgeons sometimes encounter situations where reaching hemostasis is difficult, including disseminated intravascular coagulation owing to cancer, pregnancy, infection, massive tissue injury, and uncontrolled oozing from needle holes under heparinization in cardiovascular surgeries. There are a wide variety of hemostatic agents with various types of materials and processes. The basic mechanism of most commercially available agents is to accelerate blood coagulation via fibrin formation and platelet activity. However, the hemostatic effects can be negatively affected by abnormal coagulation conditions. Synthetic sealants using polymerization have been developed under strong demand for reliable intraoperative hemostatic agents independent of blood conditions, especially in the field of cardiovascular surgery (Fig 1).

**Fig 1.**
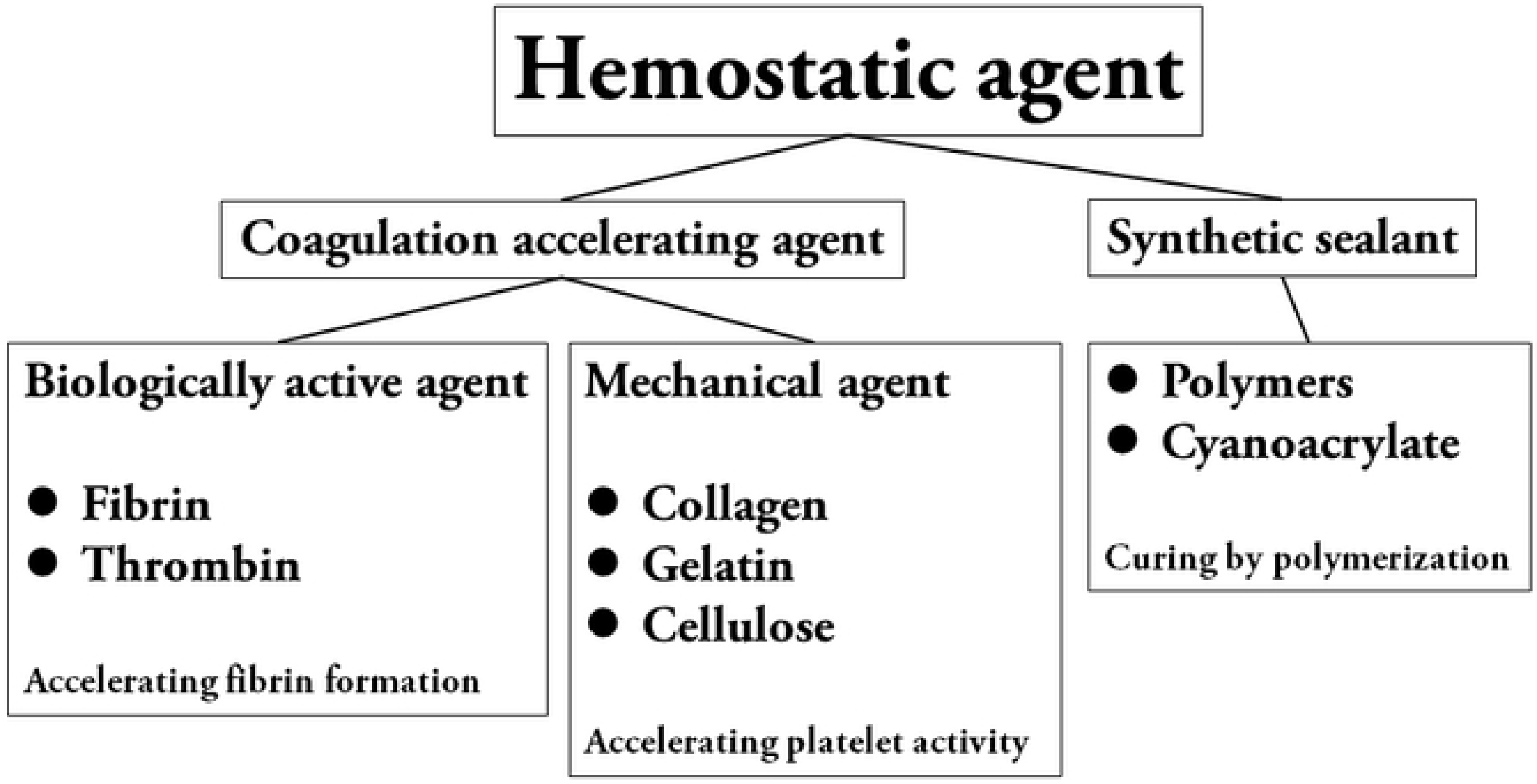
Categorization of hemostatic agents.

We designed a next-generation synthetic sealant using a polyethylene glycol (PEG) hydrogel (TetraStat) that solidified instantly in response to pH changes. Formed by the reaction of two types of tetra-armed PEGS, TetraStat responds slowly in acidic conditions and extremely fast in neutral conditions [1,2]. When TetraStat comes in contact with a neutral body fluid such as blood, apparent solidification occurs. This principle, which is independent of the biological blood coagulation reaction, allows for the selective chemical sealing of bleeding sites. Therefore, this non-biomaterial sealant could be used in situations with uncontrolled hemorrhaging and should be theoretically effective under anti-coagulant and anti-platelet drug administration and abnormal coagulation states.

In this study, we evaluated the ex vivo and in vivo effectiveness of TetraStat. First, we applied TetraStat to various sized holes in a prosthetic graft connected to a circulatory system filled with phosphate-buffered saline (PBS) under systemic pressure. Next, we used a porous PEG sponge as a temporary liquid reservoir for TetraStat and applied it to a large hole in the vena cava of a rat.

## Methods

### Preparation of TetraStat

We used mutually reactive tetra-armed PEGs with sulfhydryl groups at the termini (Tetra-PEG-SH; SUNBRIGHT PTE-100SH, NOF CORPORATION, Tokyo, Japan) and maleimide groups at the termini (Tetra-PEG-MA; SUNBRIGHT® PTE-100MA, NOF CORPORATION, Tokyo, Japan). Aqueous solutions of tetra-PEG-SH and tetra-PEG-MA were prepared by dissolving the respective PEG in 1 mM citrate phosphate buffer (pH 3.0) at the desired polymer concentrations for each experiment. Equal volumes of tetra-PEG-SH and tetra-PEG-MA solution were combined to form the TetraStat solution. A porous PEG sponge (Gellycle Co., Ltd., Tokyo, Japan) was then soaked in an excess amount of TetraStat.

### Ex vivo experiments

A circulatory system was created using a silicone tube connected to a 6-mm diameter expanded polytetrafluoroethylene (ePTFE) prosthetic graft (Propaten, Gore Medical, Flagstaff, AZ, USA). This closed circuit was filled with PBS under human systemic pressure (100-150 mmHg), which was controlled and measured using a liquid fluid pump and pressure gauge (Fig 2).

**Fig 2.**
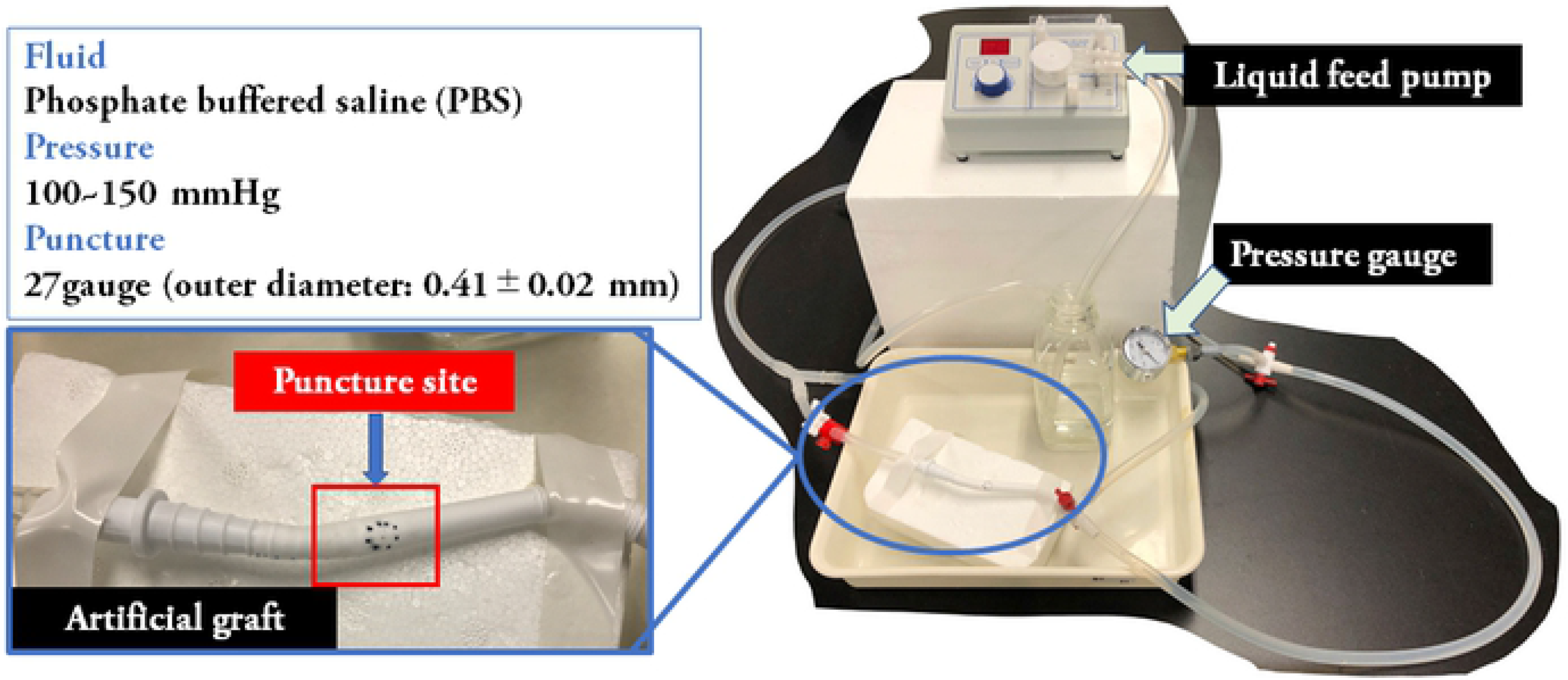
Ex vivo circulatory system.

We punctured the graft using a 27-gauge needle (outer diameter: 0.41±0.02 mm). A porous PEG sponge soaked with TetraStat was applied to the punctures and compressed for 1 min. To compare the hemostatic capability of TetraStat with that of existing products, we chose two products widely used in Japan: oxidized cellulose (SURGICEL, Johnson & Johnson Medical Ltd., Tokyo, Japan) and a fibrinogen and thrombin sealant patch (TachoSil, Takeda Nederland b.v. Takeda, Zurich, Switzerland).

### In vivo experiments with rats

All animal experiments were performed according to the Guideline for the Care and Use of Laboratory Animals established by the University of Tokyo (Permit Number: KA20-7, approval data; August 11, 2020). Male Sprague Dawley rats (320-380 g of body weight) were used for the experiments. They were fed a normal diet and were kept in air conditioning (21±1°C) with a 12-h light-dark cycle.

The rats were anesthetized using isoflurane. Laparotomy was performed at the midline, and the infrarenal vena cava was exposed. Each vena cava was punctured with a different size needle (14, 18, or 20 gauge) 1 cm above the junction of the left and right common iliac veins. After withdrawing the needle, bleeding occurred for 5–7 s, and the punctures were then astricted with the three hemostatic agents for 1 min. To adhere to the commercial instructions for use, SURGICEL and TachoSil were also applied for 5 min.

### In vivo evaluation of hemostasis

We removed any pooled blood and confirmed hemostasis macroscopically. Then, the abdominal incision was closed, and the subject was recovered from anesthesia. Seven days after the operation, the rats were euthanized with an isoflurane overdose. A 2-cm section of the treated vena cava was excised and fixed in a buffered 4% formalin solution for 24 h. After formalin fixation, the excised vessels were embedded in paraffin. The embedded tissue was cut into 5-mm thick slices and stained with hematoxylin and eosin (HE) stain. The inflammatory reaction was evaluated using a grading criterion from 0 to 3 (0, no inflammation; 1, mild inflammation; 2, moderate inflammation; and 3, large amounts of inflammation) [3].

## Results

### Ex vivo experiments

The punctures were successfully sealed in most TetraStat cases (S1 Video). In one case with 50 g/L TetraStat, the sealing was unsuccessful. However, with concentrations ranging from 60 to 100 g/L, we increased the intraluminal pressure up to 600 mmHg after sealing was confirmed without leakage (Table 1). Ex vivo sealing was unsuccessful in cases in which SURGICEL and TachoSil were used.

**Table 1.**
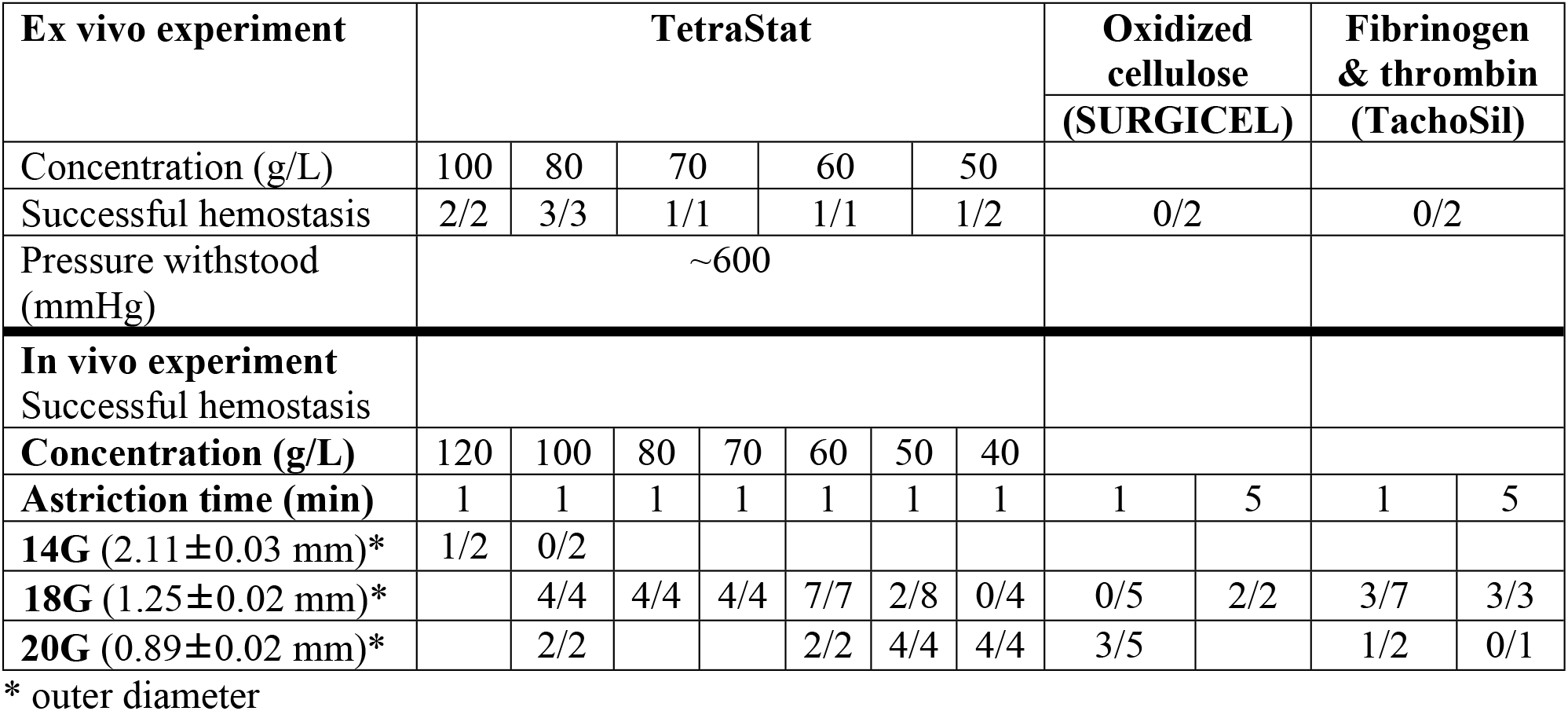
Results of sealing the punctured sites (“hemostasis”).

### In vivo experiments

During the in vivo experiments, most punctures successfully sealed after 1 min astriction with TetraStat, despite needle rotation and allowing bleeding for several seconds with subsequent massive bleeding (S2 Video, Fig 3). Using the 20-gauge needle, all applied TetraStat concentrations were effective. Successful hemostasis was also observed in some cases in which SURGICEL and TachoSil were used. For the 18-gauge needle with an outer diameter approximately the same as the vena cava diameter, hemostasis seemed less effective with 40 and 50 g/L TetraStat, indicating a dose-dependent effect. In the SURGICEL and TachoSil groups, we astricted and crushed the vena cava for 5 min and found successful hemostasis after puncture with an 18-gauge needle. Hemostasis was also successful following puncture with a 14-gauge needle in one case with 120 g/L TetraStat (Table 1).

**Fig 3.**
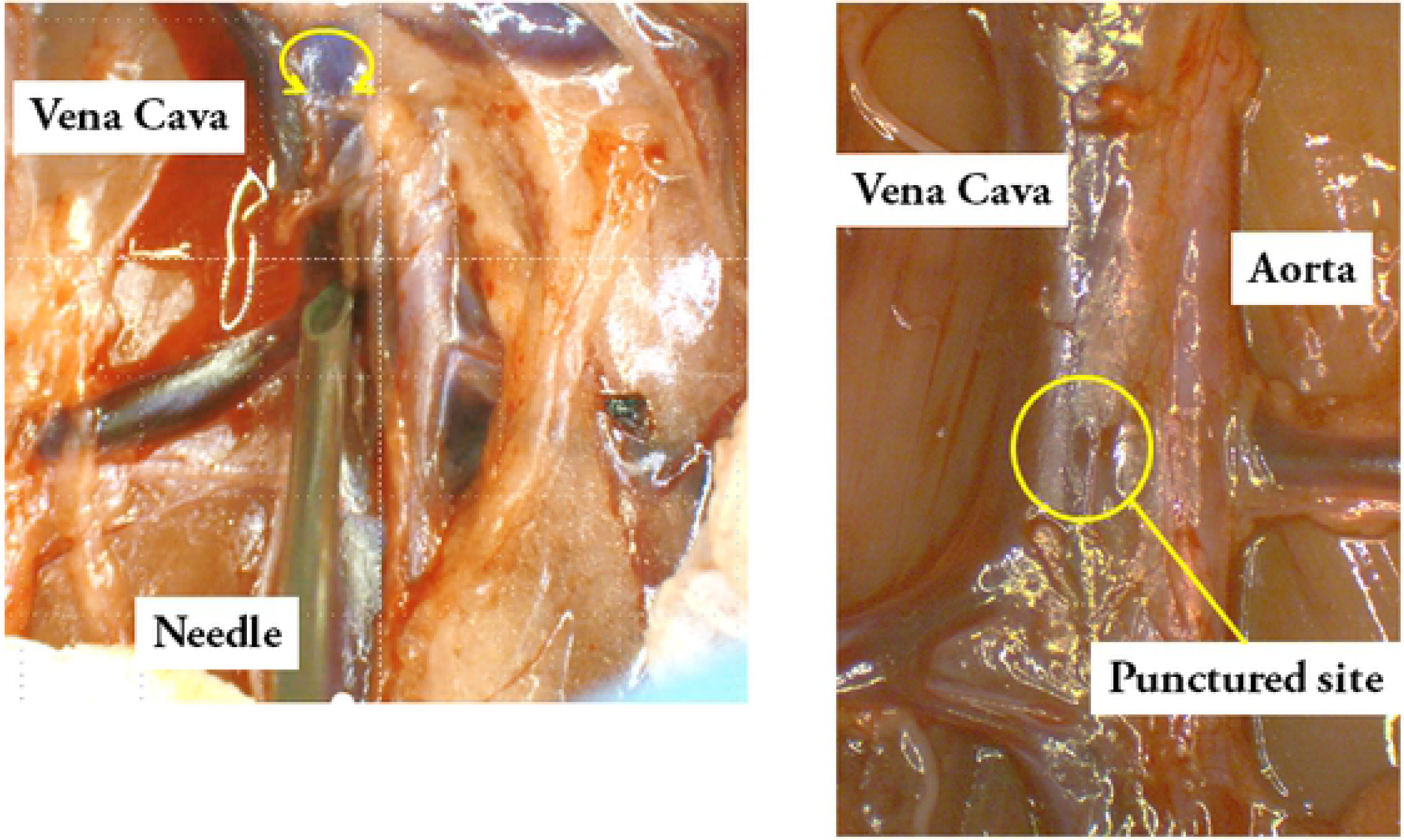
In vivo model of a needle puncture in the vena cava of a rat.

### Adverse events

Because of the acidity of TetraStat, postoperative intraperitoneal inflammation was expected. Severe adhesion was found in 2 out of 8 and 1 out of 4 cases with 100 g/L and 80 g/L TetraStat, respectively. However, there were no macroscopic adhesions below 80 g/L TetraStat.

### Histological findings

HE-stained sections from the puncture sites sealed with TetraStat, SURGICEL and TachoSil (n=4 for each group) were investigated and compared. The amount of residual agent on the puncture site was significantly lower in the TetraStat group (Fig 4A). Inflammatory reactions, including inflammatory cell infiltration, granulation, and fibrosis, were found in all groups; grade 1 reactions were seen with TetraStat, and grade 3 reactions were seen with SURGICEL and TachoSil.^3^ Remarkable venous wall thickening with severe inflammatory cell infiltration was found in the SURGICEL group (Fig 4B). Thrombi in the vena cava were found in the SURGICEL group (1 case) and the TachoSil group (3 cases) but were not found in the TetraStat group. (Fig 4C)

**Fig 4.**
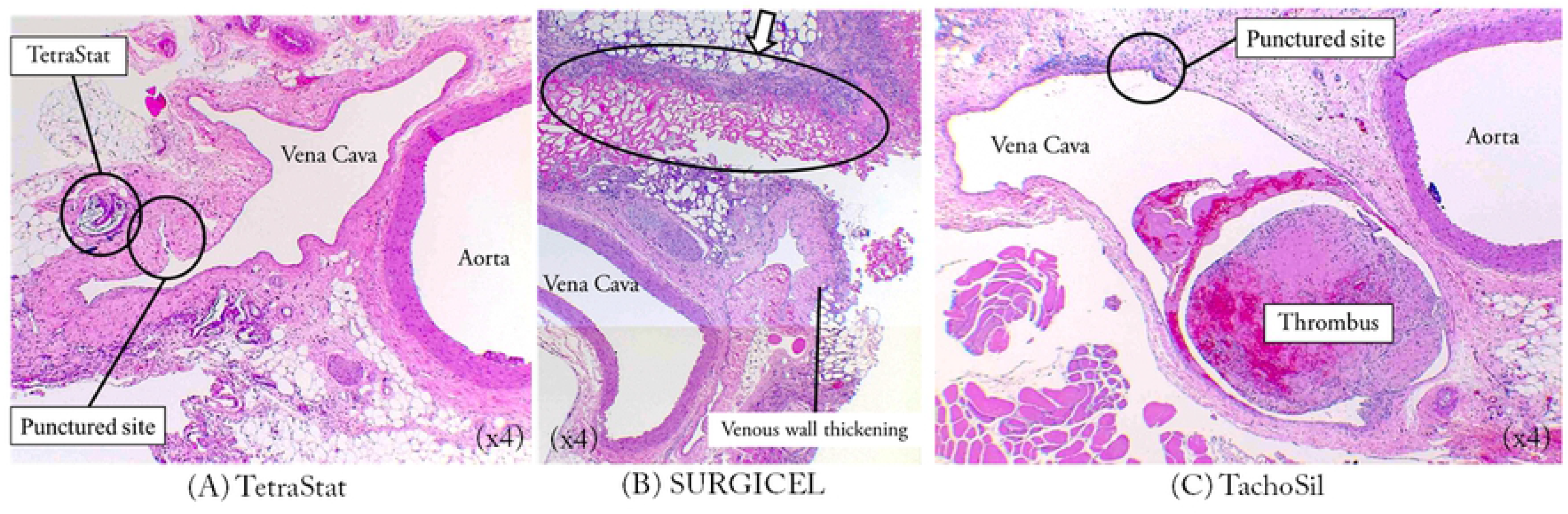
Histological findings (HE staining). (A) TetraStat. Small amount of the remnant agent with mild inflammation is seen near the punctured site. (B) SURGICEL. Significant residual agent with severe inflammation is seen along the wall of vena cava. The wall thickens remarkably, accompanied with inflammatory cell infiltration. (C) TachoSil. A thrombus is seen in the vena cava.

## Discussion

We demonstrated the high hemostatic ability of TetraStat in both ex vivo and in vivo experiments. For comparison, we selected two commercially available hemostatic agents that are widely used in Japan. TachoSil is a reliable biological sealant with high hemostatic ability but is expensive owing to limited availability. SURGICEL, while popular because of its low cost and non-biological nature, is usually used only supportively after suture hemostasis because of its relatively low hemostatic ability. Initially, we focused on the synthetic nature of TetraStat, which promised no risk of infection from biological pathogens and hemostatic mechanisms independent of the blood coagulation process and controllable gelation. However, TetraStat’s sealing ability was unexpectedly high. Inflammation occurred after intraperitoneal administration of high concentrations of TetraStat, most likely because of its acidity. However, SURGICEL, which has been widely used for several decades, also has a high acidity [4]. Given its high hemostatic performance, inflammation with TetraStat might be an acceptable clinical adverse event.

The use of the porous PEG sponge was the key to this hemostatic system, which was effective even under massive hemorrhage. Initially, we attempted to only use the solidifying liquid. However, we found that the liquid became diluted, and a large amount of TetraStat was needed for massive hemorrhages occurring under blood pressure. We overcame this issue by employing the porous PEG sponge as a temporary liquid reservoir for the TetraStat, thereby preventing the liquid from mixing with or becoming diluted by the blood. Furthermore, the blood-TetraStat composite was removable after hemostasis confirmation (Fig 5).

**Fig 5.**
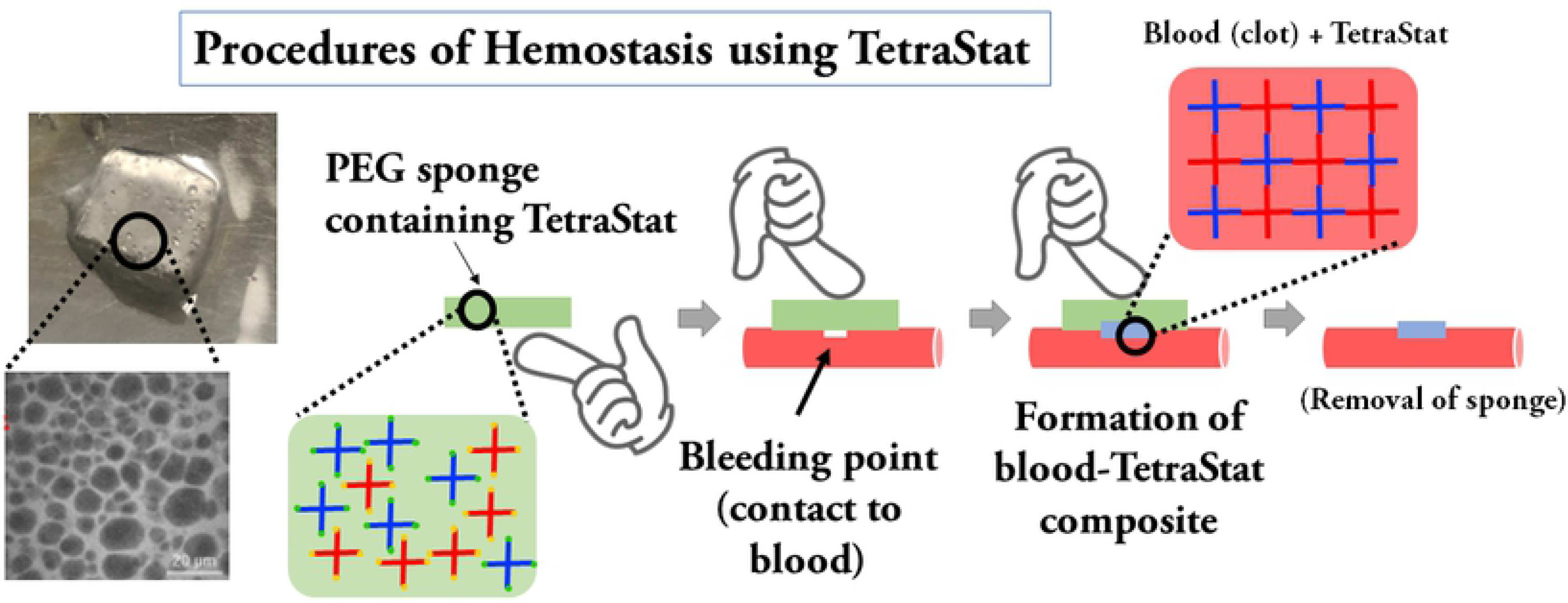
Hemostatic system using a porous PEG sponge soaked with TetraStat.

Hydrofit (Sanyo Medical Industry, Ltd., Kyoto, Japan) is a novel synthetic sealant recently developed in Japan and formed by a reaction between a copolymer of PEG and fluorinated hexamethylene diisocyanate (FHD). Hydrofit has a high hemostatic ability and is safe and effective in thoracic aortic surgery [5]. However, this liquid agent can be difficult to use, thus requiring skill and experience [6]. In addition, the long-term safety of residual FHD after hemostasis has not been confirmed. The TetraStat hemostatic system using a PEG sponge was used in massive hemorrhage and then removed after hemostasis. After removal of the gel, a thin PEG component was found at the puncture site histologically; theoretically, the PEG could be adjusted to be biodegradable [7]. Fortunately, intraluminal thrombosis was not found in any specimens, possibly because of the sponge’s structure.

An attempt to use PEG-based hydrogels for rabbit liver injuries was previously reported by Bu et al [8]. The hydrogel liquid was directly applied to the injured liver, and gelation occurred after a certain period of time. While potentially useful with very small amounts of bleeding or for postoperative bleeding, their hydrogel material did not exhibit the blood-responsive gelation observed with the TetraStat and PEG sponge system. Therefore, ongoing bleeding may dilute the hydrogel liquid, rendering the material ineffective. In addition, a porous sponge is assumed clinically necessary for pinpoint hemostasis in massive bleeding lesions with high blood pressures. With our design, the hemostatic component is not easily diluted with blood during massive bleeding. These factors may contribute to the cost-effectiveness.

In our in vivo experiments, we injured the vena cava and not the aorta. Although the aorta was an appropriate site for puncture owing to its high blood pressure, the difference in hemostatic effect between the agents was not remarkable because of the thickness of the aortic wall. In addition, the demand for sealants for uncontrollable venous hemorrhage is high because venous injury repair is considered technically difficult and requires experience owing to the thin fragile venous wall structure. We successfully created a simulation in rats mimicking the uncontrollable hemorrhage seen by clinical surgeons.

Unlike in clinical cardiovascular situations, we did not perform heparinization in the in vivo experiment. Nevertheless, we demonstrated the impact of TetraStat compared to commonly used hemostatic agents. Our next step is to use larger animals to confirm the safety and effectiveness in models more closely mimicking humans. For clinical use, optimizing the concentration of the sealant is necessary.

In conclusion, ex vivo and in vivo experiments revealed the high hemostatic ability of TetraStat to seal punctures under high pressure and large injuries in the rat vena cava during massive hemorrhage. A porous sponge of PEG soaked with TetraStat is a promising advancement in hemostatic systems.

## Acknowledgments

The porous PEG sponge used in the experiments was a generous gift from Gellycle Co.

## Supporting information

S1 Video. Demonstration of the successful use of TetraStat.

S2 Video. Demonstration of the successful use of TetraStat.

## Notes

### Competing Interest Statement

The authors have declared no competing interest.

